# Computational vaccinology based development of multi-epitope subunit vaccine for protection against the Norovirus’ infections

**DOI:** 10.1101/2020.03.18.997197

**Authors:** Irfan Ahmad, Syed Shujait Ali, Ismail Shah, Shahzeb Khan, Mazhar Khan, Saif Ullah, Shahid Ali, Jafar Khan, Mohammad Ali, Abbas Khan, Dong-Qing Wei

## Abstract

Human Norovirus belong to family *Calciviridae*, it was identified in the outbreak of gastroenteritis in Norwalk, due to its seasonal prevalence known as “winter vomiting disease”. Treatment of Norovirus infection is still mysterious because there is no effective antiviral drugs or vaccine developed to protect against the infection, to eradicate the infection an effective vaccine should be developed. In this study capsid protein (A7YK10), small protein (A7YK11) and polyprotein (A7YK09) were utilized. These proteins were subjected to B and T cell epitopes prediction by using reliable immunoinformatics tools. The antigenic and non-allergenic epitopes were selected for subunit vaccine, which can activate cellular and humoral immune responses. Linkers joined these epitopes together. The vaccine structure was modelled and validated by using Errat, ProSA and rampage servers. The modelled vaccine was docked with TLR-7. Stability of the docked complex was evaluated by MD simulation. In order to apply the concept in a wet lab, the reverse translated vaccine sequence was cloned in pET28a (+). The vaccine developed in this study requires experimental validation to ensure its effectiveness against the disease.

## Introduction

Norovirus (NVs), also known as “small round-structured virus” and this RNA single-stranded virus is placed within family *Calciviridae.* NVs is responsible for the spread of non-bacterial human gastroenteritis (1). The Human norovirus (HuNV) was first identified in stool specimen, during gastroenteritis outbreak in Norwalk and was coined as Norwalk virus. More than a thousand strains of NVs are isolated which are genetically and serologically different. The infected person has an abdominal cramp, stomach pain, diarrhoea, vomiting and nausea with mild pyrexia (2). The consumption of contaminated food and water is deemed essential for the development and spread of disease (3), (4), and globally 20 % of all diarrheal diseases are caused by HuNV, and nearly 21200 victims succumbed to death annually (5, 6). HuNV interferes with interferon type I & III by influencing MHC-I expression and causing rapid infection. MHC-I plays a key role in providing immunity against viruses. In this process, proteasome-mediated degraded peptides are presented to the CD8+ T-cells for evoking immune reactions (7). The genome of HuNV is 7.5 kb, which consists of three open reading frames (ORF’s), ORF1, ORF2, and ORF3. These ORF’s (ORF1, ORF2, and ORF3) codes for a nonstructural protein, VP1 major capsid protein, and VP2 minor capsid proteins respectively (8).

In clinical samples, an electron microscope (EM) (9) is used as a diagnostic tool for norovirus identification. ELISA (enzyme-linked immunosorbent assay) and molecular techniques are accessible for the diagnostic purpose of pathogens including norovirus. EM is found in the well-equipped lab and it is used to look for the pathogenic particles in the feces. RT-PCR (10) shows sensitivity for identification, and it also assist in comprehension of these viruses molecular diversity (11-15). Genomic characterization and molecular diversity is assessed by (HMA) hetero duplex mobility assay of various viruses which include; Norovirus (16), measles virus (17), polioviruses (18), hepatitis C virus (19) and polioviruses (18). Cross challenge studies and IEM (immune electron microscopy) studies (20) was previously utilize for NV antigenic diversity assessment, before the development of rNV (recombinant-NV) capsid protein.

Non-bacterial gastroenteritis is still a great challenge, and there is no effective licensed vaccine available for its treatment (7). Researchers are trying their best to launch an effective vaccine against norovirus, however, their investigations are either in clinical trials or in pre-clinical stages (21). These investigations may results in two norovirus vaccines in future, bivalent GI.1/GII.4 intramuscular VLP vaccines (in phase II b clinical trial), and monovalent GI.1 oral pill recombinant adenovirus vaccine (phase I trials) (7). NoV infection is equally pervasive in developing as well as developed countries. Children and elders are severely affected by these infections. Effective vaccines and drug designing would be instrumental in controlling the higher mortality rates caused by these infections (6). Vaccines evoke innate and cellular immune responses to develop antibodies and memory cells, which may provide long-lasting protection from specific serotypes (22). In-silico study based on immunoinformatics approaches was applied to pinpoint effective epitopes or hits as a potential candidate for vaccine or drug designing (23-27). Computational approaches are beneficial to predict the antigen without culturing the pathogenic strain experimentally (28-32).

In this study, computational analysis was performed to predict the epitopes based effective vaccine against the NoV. T-cell, B-cell and HTL epitopes were predicted and analyzed using defined criteria for selecting the potential epitopes for final vaccine construct. Further implementation of molecular docking with TLR receptors, molecular dynamics simulation and codon optimization for expression confirmed the potential of the final vaccine construct. Thus, this study provides a way towards the development of a potential vaccine candidate against the Norovirus.

## Results

### Protein collection

The amino acid sequences of capsid protein (UniProt id: A7YK10), polyprotein (UniProt id: A7YK09) and small protein (A7YK11) of *Norovirus* as well as Beta-defensin 3 (Q5U7J2), an adjuvant was retrieved from UniProtKB in FASTA format. These protein were antigenic in nature based on antigenicity score of 0.48, 0.52, and 0.49 respectively as calculated by VaxiJen server and were selected for the designing of a multi-epitope vaccine by immunoinformatics approach.

### CTL (Cytotoxic T Lymphocytes) epitopes prediction

NetCTL1.2 server predicted a total of 51 CTL epitopes of 9-mer in length. In these epitopes, six non-allergenic epitopes **(Table 1)** were selected for vaccine designing based on high binding affinity score. Based on the predicted scores, two epitopes were selected from A7YK09, A7YK10 and A7YK11 each.

**Table 1.** Selected CTL epitopes. All the epitopes are based on their scores.

| Proteins | Sequence ID | Peptide | Affinity | affinity rescale | Cleavage | TAP | Combine score | MHC binding Epitope |
| --- | --- | --- | --- | --- | --- | --- | --- | --- |
| A7YK10 | 469 | QSDALLIRY | 0.7635 | 3.2418 | 0.9510 | 2.8490 | 3.5269 | Yes |
|  | 91 | YLAHLSAMY | 0.5251 | 2.2294 | 0.8881 | 2.7960 | 2.5024 | Yes |
| A7YK09 | 1044 | TSSGDFLKY | 0.5906 | 2.5076 | 0.9717 | 2.9870 | 2.8027 | Yes |
|  | 1513 | MQESEFSFY | 0.5161 | 2.1913 | 0.5334 | 2.9500 | 2.4188 | Yes |
| A7YK11 | 122 | AVDWSGTRY | 0.5730 | 2.4330 | 0.9756 | 3.1630 | 2.7375 | Yes |
|  | 140 | FSGGFTPSY | 0.3257 | 1.3828 | 0.9690 | 2.6770 | 1.6620 | Yes |

### HTL (Helper T lymphocytes) prediction

MHC-II prediction module of IEDB was used for Helper T Lymphocytes (HTL) epitopes prediction for HLA-DRB1*01:01, HLA-DRB1*01:02, HLA-DRB1*01:03, HLA-DRB1*01:04, and HLA-DRB1*01:05 Human alleles.. 9 HTL epitopes with the highest binding affinity were selected. The selected epitopes are situated at position 145-159,373-387,117-131 (capsid protein), 1-15, 1631-1645, 816-830, 617-631, 1089-1103 and 193-207 (polyproteins). (Table 2 represent selected HTL epitopes).

**Table 2:**
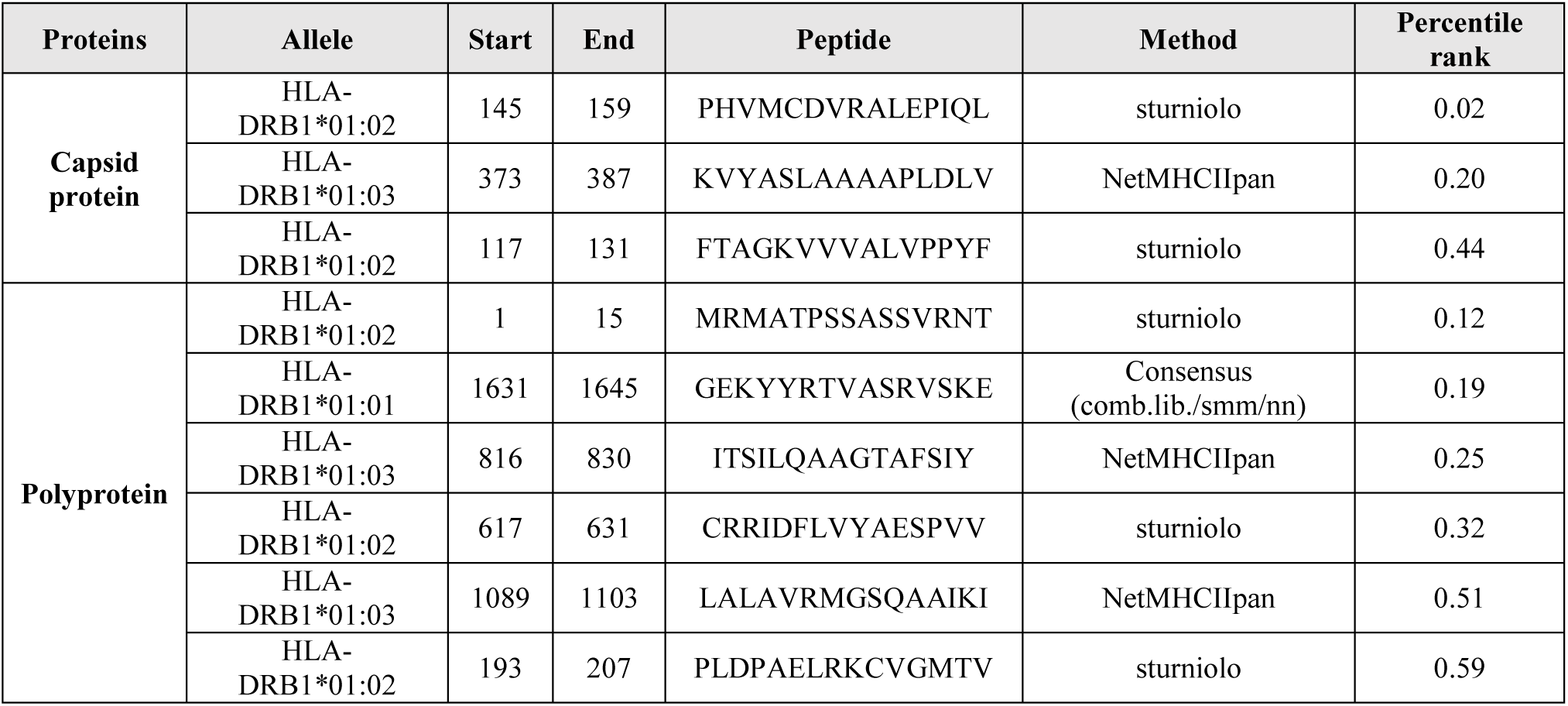
HTL selected epitopes. The epitopes are based on percentile rank.

### Interferon production

Online server IFN-epitope was used to identify the HTL epitope with a potential to induce the release of IFN-gamma from T (CD4+) cell. This analysis resulted in three epitopes (out of nine HTL epitopes) with the ability to induce T cells for interferon production **(Table 3)**.

**Table 3.**
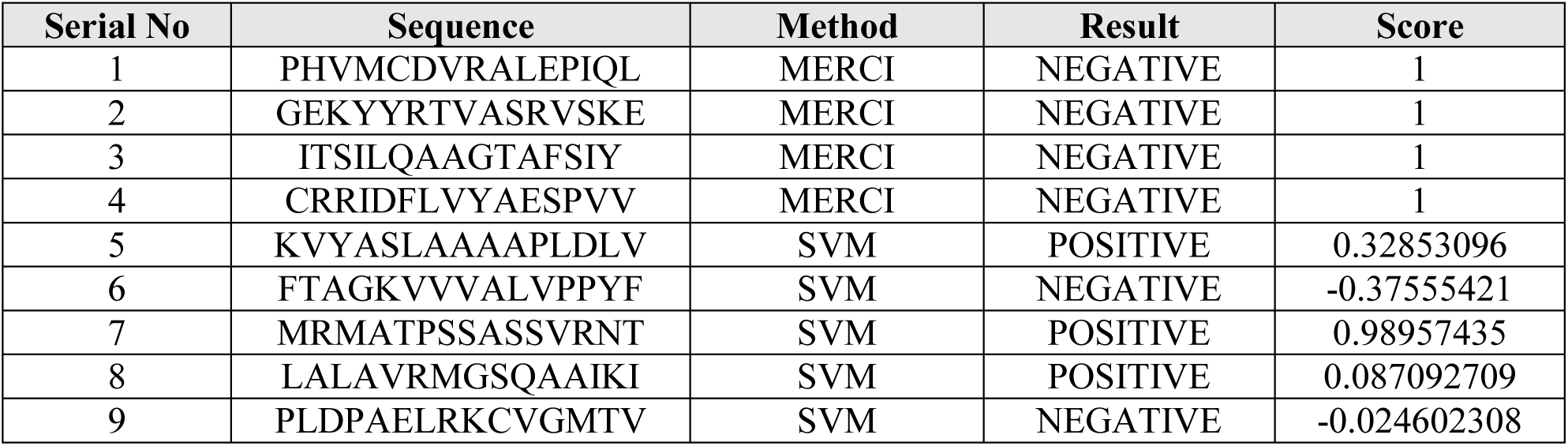
Selected HTL epitopes analysis for interferon production.

### B-cell epitope prediction for norovirus

B-cell epitope prediction was performed on ABCpreds server and six epitopes of 20-mer in length with a score higher than 0.83 were selected for further analysis **(Figure 3A)**. Conformational B-cell epitopes were identified on Discotope 2.0 and out of 359 residues, 50 B-cell residues were predicted. The predicted B-cell Linear and discontinuous epitopes are given in **Table 4** and **5**.

**Table 4:** Predicted Linear B cell epitopes with their respective scores.

| Rank | Sequence | Score | Start position |
| --- | --- | --- | --- |
| 1 | GPGGEKYYRTVASRVSKEGP | **0.91** | 222 |
| 2 | VMCDVRALEPIQLGPGPGHV | **0.89** | 127 |
| 3 | GNSISTGPGPGGGGLMGIIG | **0.87** | 334 |
| 4 | LEPIQLPLLDGPGPGMRMAT | **0.86** | 190 |
| 5 | PGPGPGVMCDVRALEPIQLP | **0.86** | 159 |
| 6 | PGPGITSILQAAGTAFSIYG | **0.84** | 241 |

**Table 5.** Predicted discontinuous B-cell epitopes residues with their contact number and Discotope score.

| S.NO | Residues | Contact number | Discotope score |
| --- | --- | --- | --- |
| 1 | LYS, TYR, PRO, GLN, ASN, GLY, PRO, GLY, GLY, GLU | 3, 6, 4, 0, 4, 1, 0, 12, 2, 4 | -2.885, -2.029, -3.589<br>-3.886, -1.879, 1.667<br>2.652, 3.693, 2.327, 0.024 |
| 2 | LYS, PRO, TYR, GLY, PRO, GLY, PRO, GLY, HIS, GLY | 8, 9, 11, 13, 3, 13, 5, 0, 0, 6 | -0.356, -2.670, -2.082,<br>1.008, 2.997, 4.536, 5.611,<br>5.290, 5.382, 5.370 |
| 3 | GLU, LYS, TYR, TYR, ARG, THR, ALA, LYS, PRO, GLY | 13, 9, 18, 32, 6, 3, 7, 0, 8, 4 | 5.690, 5.180, 4.948, 2.551, 1.623, 0.693, -2.447, -3.572, -2.696, -2.886 |
| 4 | GLY, ILE, ILE, GLY, ASN, SER, ILE, GLY, PRO, GLY | 4, 35, 7, 30, 3, 16, 3, 4, 13, 36 | -1.302, 0.705, 2.215, 3.458, 5.151, 4.587, 2.787, 2.029, 0.101, 0.667 |
| 5 | PRO, GLY, PRO, GLY, PRO, GLY, ILE, ASN, GLY, GLY | 11, 28, 7, 0, 7, 12, 6, 0, 16, 15 | -0.292, 0.722, 1.126, 1.693, 1.776, 1.315, -0.535, -2.624, 0.504, -2.062 |

**Figure. 1.**
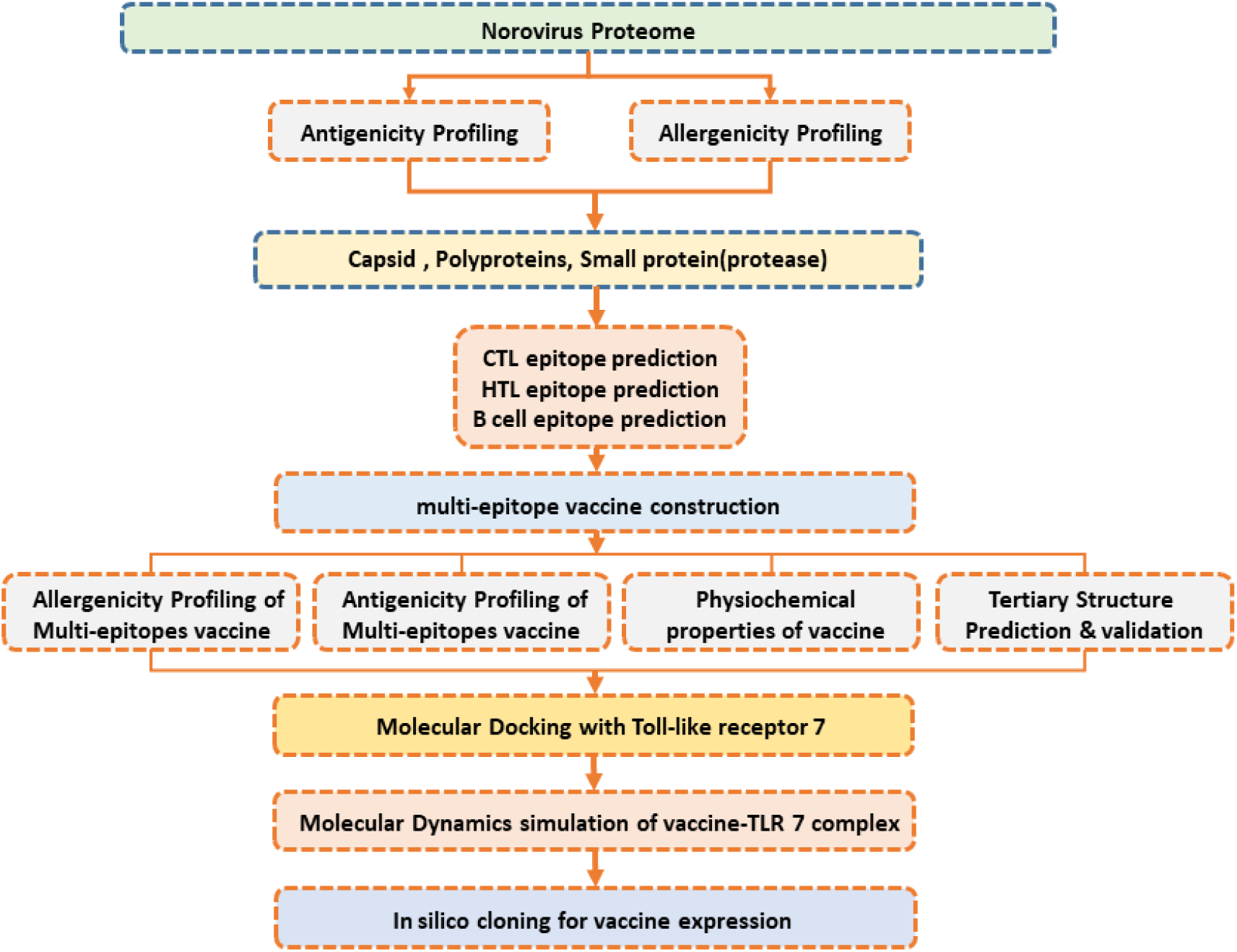
Schematic representation of the steps involved in epitope-based vaccine designing. A multi-step approach was used to construct the final vaccine candidate. Finally, validation through MD simulation and in silico expression was achieved.

**Figure. 2.**
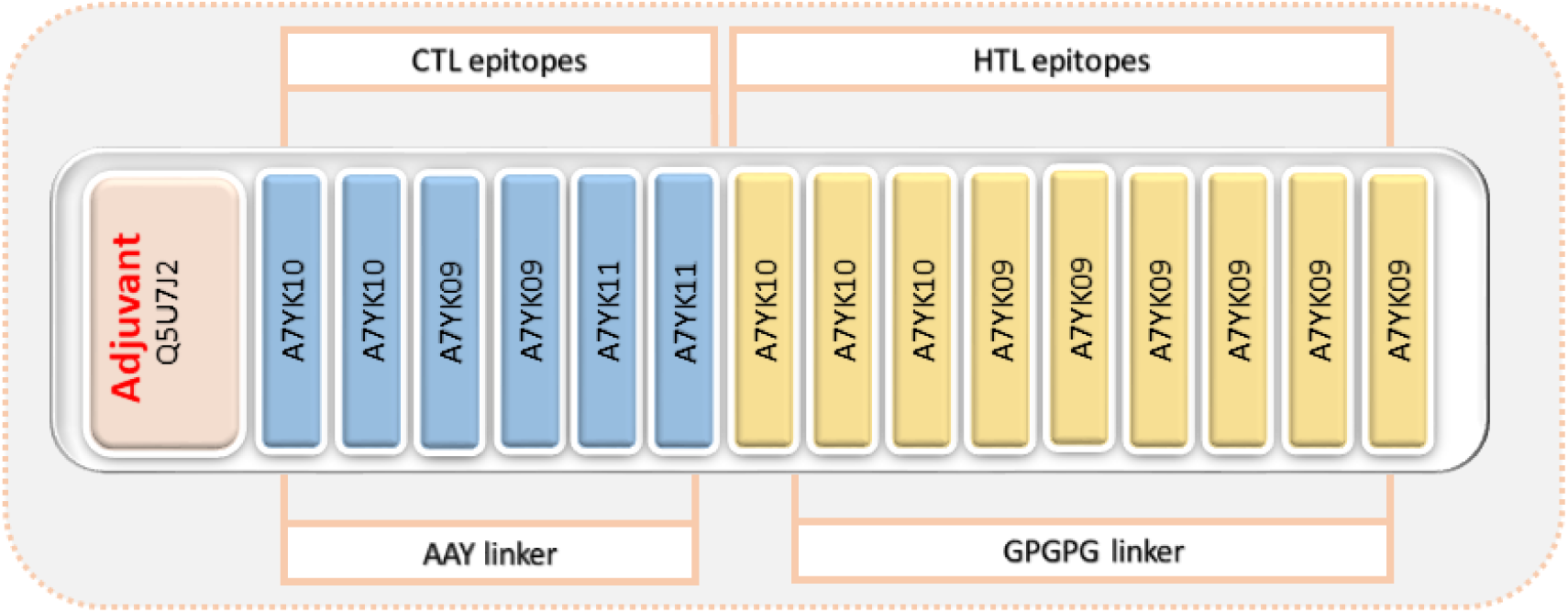
Final vaccine construct. Six different CTL epitopes while nine HTL epitopes from three different proteins were combined to construct the multi-epitope subunit vaccine using linkers. An adjuvant to the N-terminal has also been added.

**Figure 3:**
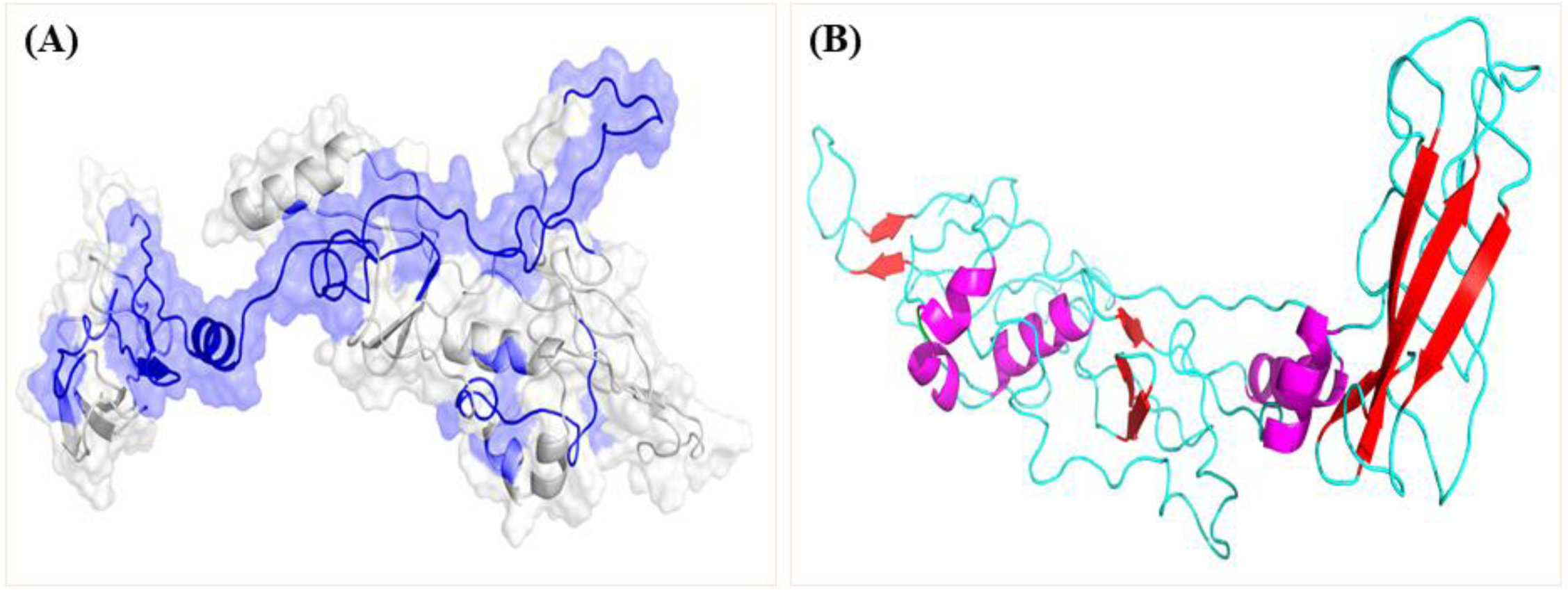
Predicted structure of the vaccine. **(A)** Showing the B-cell epitope (blue) (B) Showing the final structure of multi-epitopes vaccine construct showing loops (cyan), Helix (magenta) and beta-sheets(red).

### Final multi-epitope vaccine construct

The final multi-epitope vaccine construct was composed of 6 CTL and 9 HTL epitopes selected based on high binding affinity scores. AAY linkers were used to combine CTL epitopes and GPGPG linkers joined HTL epitopes, Whereas EAAAK linker was used for attachment of adjuvant to the N-terminal of vaccine, which amplifies its function.

### Prediction and validation of tertiary structure

3D structure of the vaccine was predicted on Robetta server. Five models were generated, and after evaluation model three (**Figure 3B**) was selected for further analysis. The selected model was validated by RAMPAGE, ProSA-web, and ERRAT (**Figure 4**). Modeled protein Ramachandran plot analysis revealed that most favored regions, additionally allowed regions, generously allowed region and disallowed region contain 81.1%, 6.8%, 1.2% and 0.4% of residues, respectively. ProSA-web and ERRAT were used for the evaluation of potential and quality errors in 3D crude model. Modeled protein overall quality factor was found 69.2% utilizing ERRAT. ProSA-web is utilized for prediction of Z-score prediction, which is found as - 4.7.

**Figure 4:**
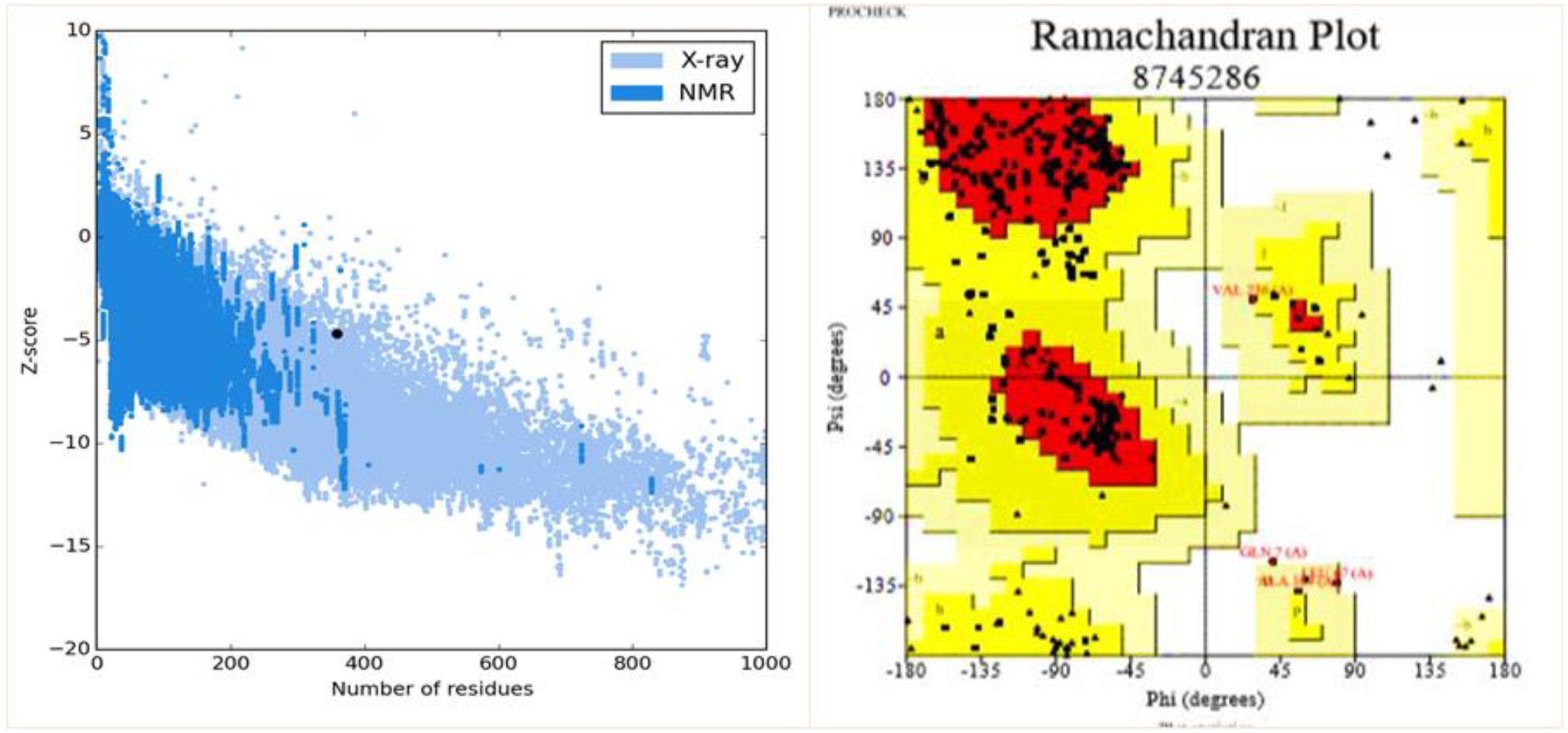
Validation of 3D final vaccine model. **(A)** PROSA showing Z-score (−4.7) for 3D structure validation **(B)** In Ramachandran analysis residues were allocated; most favoured region 81.1%, allowed 6.8%, generously allowed 1.2% and disallowed region 0.4% residues.

### Antigenicity, allergenicity and physiochemical parameter prediction of final vaccine construct

The antigenicity of the final vaccine construct was predicted on VaxiJen and ANTIGENpro servers by selecting bacteria model at 0.4% threshold. The antigenicity scores predicted by VaxiJen and ANTIGENpro are 0.8134% and 0.635949%, respectively, which indicates the antigenic nature of the final vaccine. For allergenicity, AlgPred was used and −0.62811 score indicating the non-allergen nature of vaccine using the default threshold of −0.40. The physiochemical parameters including molecular weight (MW) and theoretical isoelectric point value (PI) of vaccine were found 36.56 kDa and 9.17, respectively, as predicted by ProtParam. The PI value (9.17) suggesting the basic nature of the vaccine. Moreover, the half-life in mammalian reticulocytes, yeast, and *E. Coli* was found 30 hours (in vitro); 20 hours (in vivo), and 10 hours (in vivo), respectively. The instability index score of 34.03 refers to the stable nature of the protein. The GRAVY (Grand average of hydropathicity) and the aliphatic index was found 75.88 and −0.040, respectively.

### Prediction of secondary structure

Secondary structure for vaccine predicted by PSIPRED program suggests the presence of 20.89% alpha-helix, 16.71% beta-strand, and 62.39 % coil as shown in **Figure 5**.

**Figure 5:**
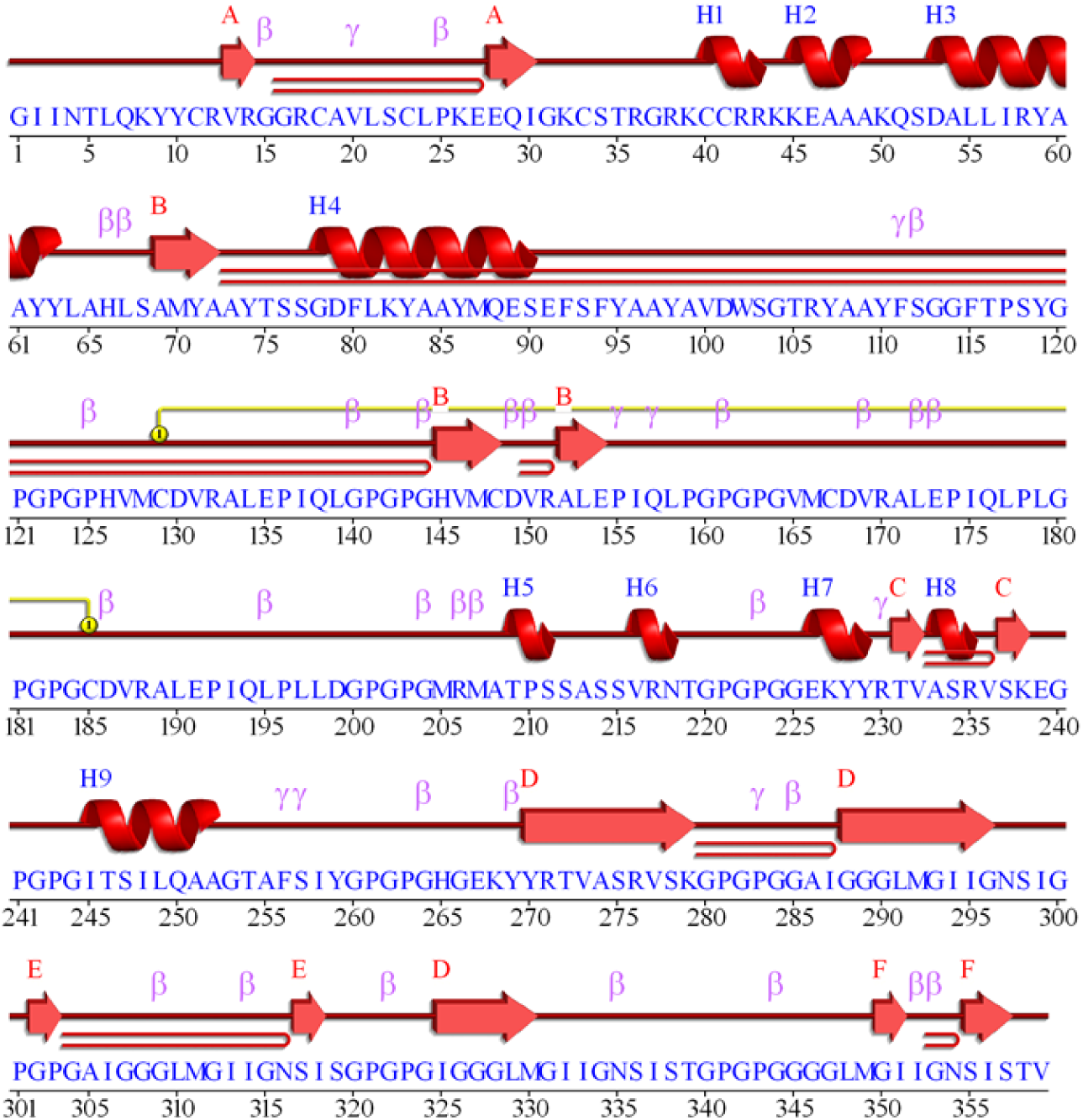
Represent the secondary structure of the final vaccine construct in which alpha helix, beta-strands and coils were identified (20.89%, 16.71% and 62.39%).

### Subunit vaccine molecular docking with an immune receptor (TLR-7)

For docking of TLR-7 was docked with the multi-epitopes vaccine, using an online server ZDOCK. Overall, ten complexes were generated and the most suitable vaccine-TLR complex was selected based on correct conformation and binding **(Figure 6)**. The PDBsum server reported 90 interface residues with one salt bridge and eight hydrogen bonds were reported between the vaccine and TLR-7.

**Figure 6:**
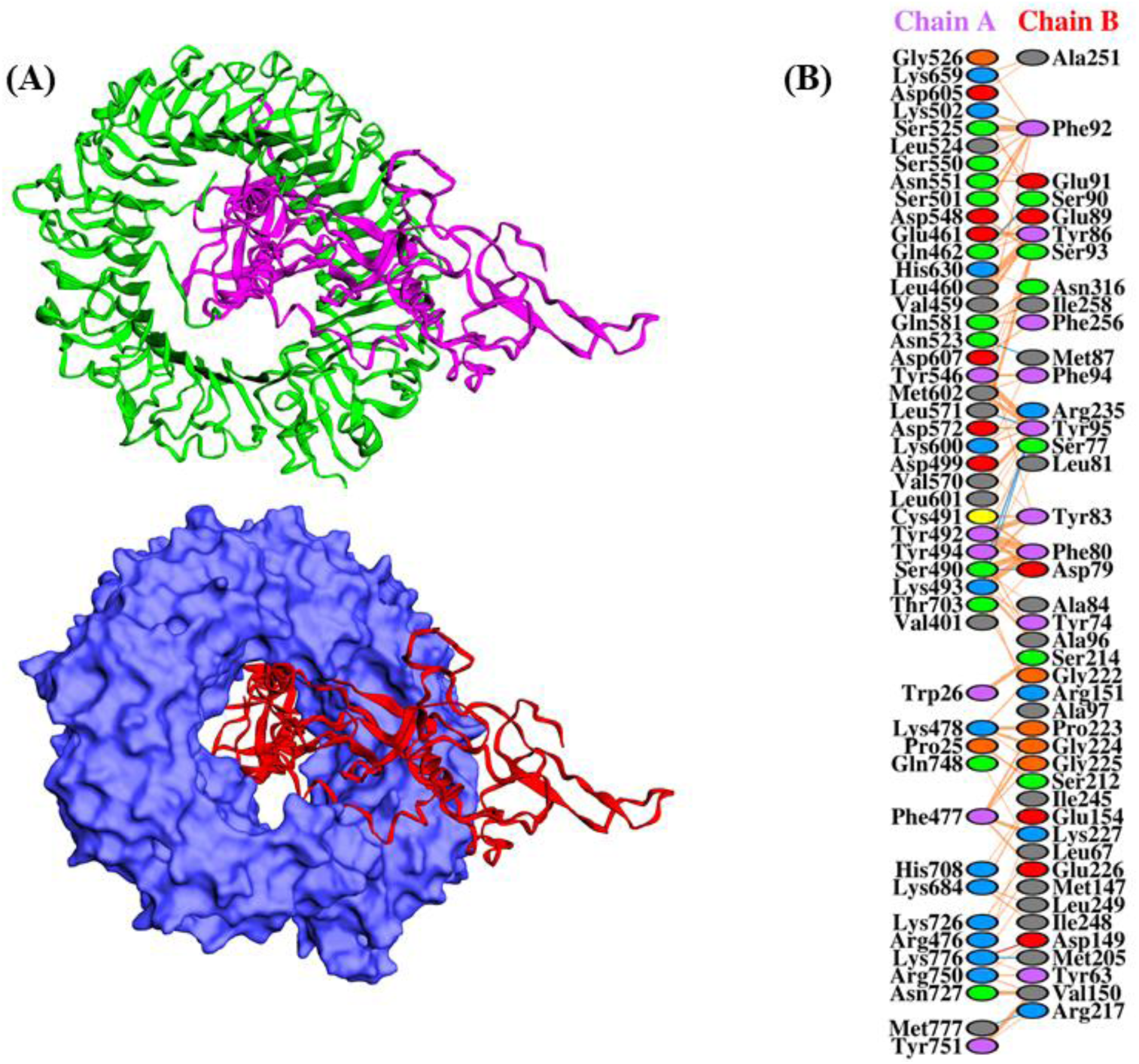
**(A)** TLR-7 (PDB ID: 5GMF) and vaccine docked complex. Magenta colour represents the vaccine while the green and blue (surface) colour represents receptor. **(B)** Showing the interaction pattern of the TLR-7 and vaccine construct.

### MD simulation of immune receptor-vaccine complex

Molecular dynamics (MD) simulation was performed to check the stability and fluctuation of vaccine construct and TLR-7 complex. The computed RMSD and RMSF for the vaccine (protein) and its side chain as well as their graph are shown in **Figure 7**. The RMSD and RMSF of protein and side-chain residues respectively were checked at 30ns time to estimate the stability of the system. Overall fluctuation (RMSD) rate for the simulated system was found 4Å (TLR-7) and RMSF (residual fluctuation) for maximum residues were found in acceptable range while some residues exhibit higher fluctuation.

**Figure 7:**
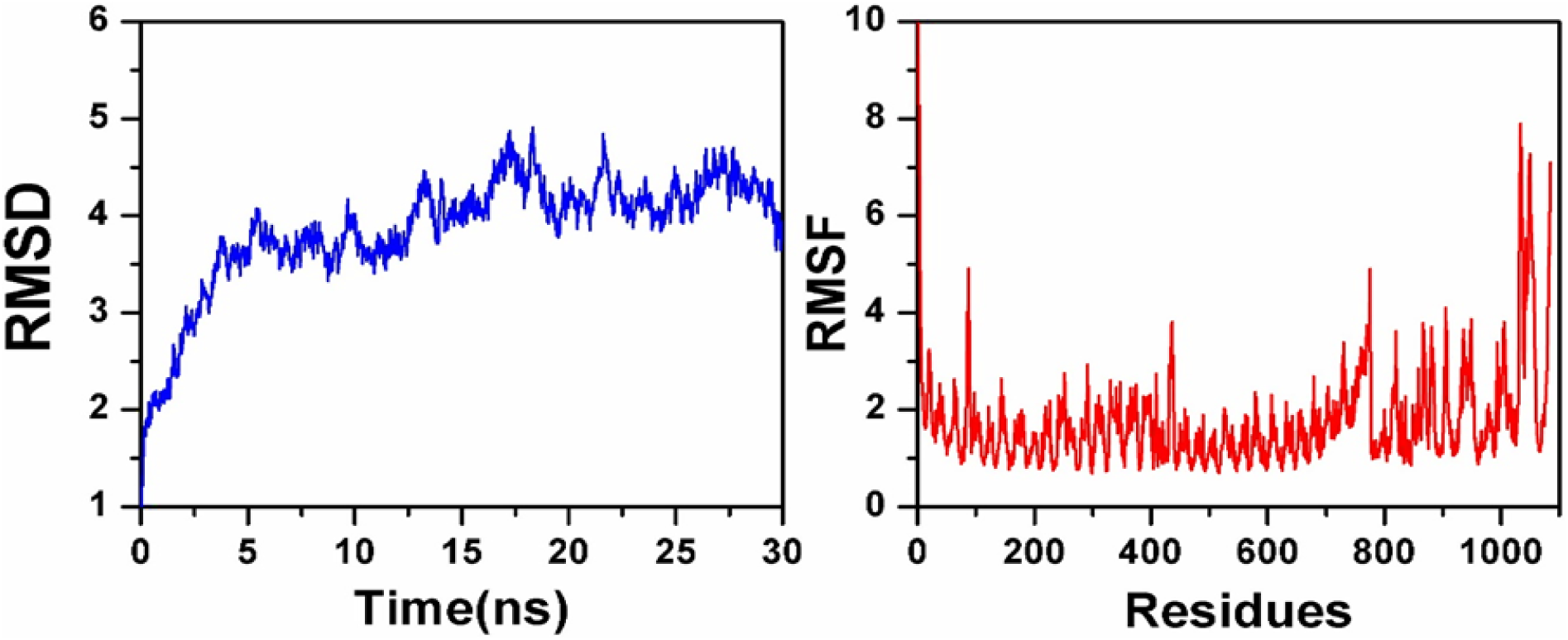
Molecular dynamics simulation of the receptor-vaccine complex. The left graph is showing the RMSD of the complex (X-axis = Time in ns and Y-axis = RMSD) while the right graph is showing the RMSF of the complex (X-axis = Time in residue and Y-axis = RMSF).

### Codon optimization and in silico cloning

To assure the maximal expression of the protein, vaccine codon was optimized in *E. coli* (strain k12) using Java Codon Adoption Tool. The optimized length of the codon sequence is 1077 nucleotides. The average GC content was found 56.7% (optimum range 30%-70%) and CAI (codon adaption index) was 0.968, which indicates possibilities of good expression in host *E. coli*. Finally, using restriction enzymes restriction clone was formed and adapted codon sequence was inserted in pET28a (+) plasmid. The designed construct is shown in **Figure 8**.

**Figure 8:**
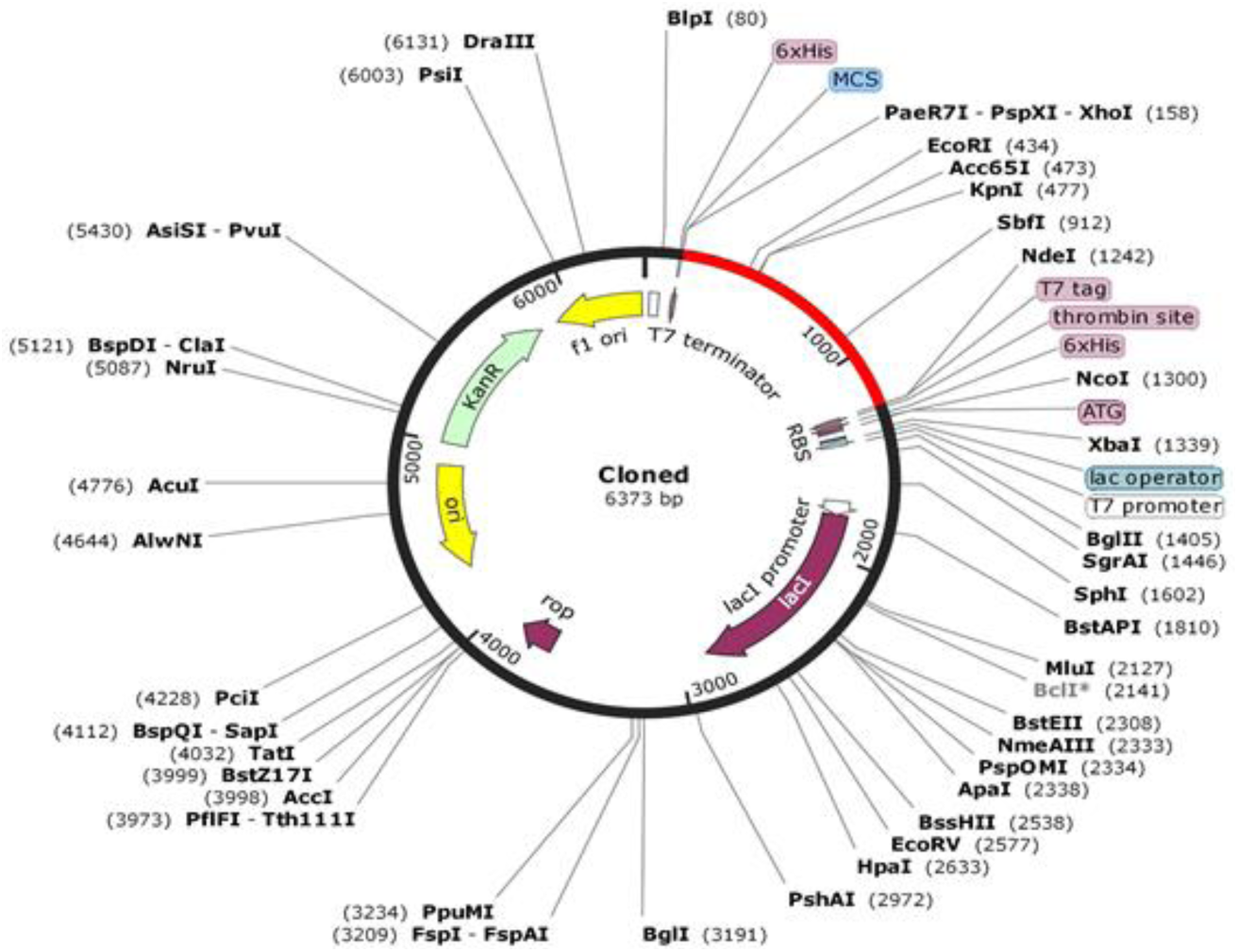
In silico restriction cloning of the final vaccine construct into pET28a (+) expression vector where Red part representing the vaccine insert and black circle showing the vector.

## Discussion

Norovirus proteins such as capsid, polyprotein, and protease were found antigenic and are vital for infection and replication within the host. Therefore, they are considered very important for subunit vaccine analysis. Immunization or vaccination is universally recognized method to eradicate or control the infection. The advancement in computational approaches and their applications to biological research ushered a new era of subunit vaccination designing in which the most accurate or exact antigenic part is identified and used as an immunization tool instead of the whole pathogen. The fast accumulation of vast genomic and proteomic data of pathogens including norovirus is helpful in the development of epitope-based effective vaccines to control and eradicate the diseases caused by various pathogens. Computational approaches predicted epitopes (CTL & HTL) based on norovirus proteins and validation scores of these epitopes suggest their use for subunit vaccine construction. MHC (major histocompatibility complex) is of different types, MHC-I carry a peptide of 9-mer to the surface of the cell and act like an impulsive signal for cytotoxic T cell. Which lead to cell destruction by activating immune complementary cascade. MHC-II molecules present peptide of 15-mer to Helper T lymphocytes. Our final subunit vaccine is composed of high-affinity CTL and HTL epitopes to elicit immunity. The allergenicity and antigenicity values of the vaccine were calculated, and these values indicate the non-allergen nature of the vaccine and are capable to provoke immune response due to antigenic nature.

In addition to these epitopes, B cell linear epitopes were also predicted which help in B-cell maturation, in order to produce antibodies. Physiochemical properties of vaccines such as molecular weight, theoretical PI, aliphatic array and thermal stability were calculated. The molecular weight of the vaccine was 36.560kDa, which is a suitable range for subunit vaccine; the theoretical PI score is 9.17, which indicates that vaccine is basic in nature. The aliphatic array suggests that vaccines have aliphatic side chains and instability index endorses the thermally stable nature of the vaccine. For prediction and analysis of the secondary structure of vaccine, PSIPRED V3.3 was used, which indicates the presence of 20.89% alpha-helix, 16.71% beta-strand, and 62.39 % coil. Besides, the 3D structure obtained by homology modeling work comprises of adequate information on the spatial arrangement of such essential protein residues as well as useful guidance in the study of protein normal function, dynamics, and interaction with ligand as well as other proteins. To pinpoint the error in the final 3D structure of vaccine different structural validation tools were used to detect errors. From the main Ramachandran plot, it was found that the overall model is satisfactory because most residues were found in the most-favour region while few were present in the disordered region.

Furthermore, the vaccine was docked with TLR-7 in order to understand the immune response towards vaccine final structure. Energy minimization was conducted to minimize the potential energy of the entire system for the overall conformational stability of the docked vaccine protein-TLR-7. Energy minimization repairs the structure’s unnecessary topology by ditching certain protein atoms and thus forms a more relatively stable structure with adequate stereochemistry is thus formed.

To obtain maximum expression (transcription and translation) of vaccine protein in the host (*E. coli* strain k-12), codon optimization was accomplished by CAI (codon adaptation index). The solubility of overexpressed recombinant protein in the host (*E. coli*) is one of the crucial requirements of many biochemical processes. Our vaccine protein shows a suitable proportion of solubility in the host. The essential goals of many mechanical and biomedical applications are to strengthen the protein. The newly designed vaccine was very effective and showing good immunogenic response in animals, however, when these vaccines were applied to humans the response was not similar due to the complexity of human immunopathology. In this study, modern immunoinformatics approaches were incorporated to develop new thermostable, cheap and effective subunit vaccines. These vaccines are safe and immunogenic to exploit the immune system to provide protection from norovirus infection.

## Conclusion

In this study, the main focus was to apply, *in-silico* approaches to design an effective multi-epitope vaccine against norovirus based on three proteins due to their antigenic nature. To design a vaccine, B and T cell epitopes were predicted, which is due to the presentation of pathogen epitope by MHC-I and II. Suitable linkers were used to fuse these epitopes. Vaccine tertiary structure was predicted and validated to ensure the functionality of the vaccine. The physicochemical properties such as antigenicity, allergenicity, and stability were computed. The vaccine was docked with TLR-7 to check the vaccine affinity towards the receptor.

Finally, for codon optimization, the protein was reversely translated which ensure maximum expression of the vaccine in the host (*E. coli*). A wet lab experimental validation is needed to assure the activity of constructed vaccine. This study can help in controlling norovirus infection.

## Methodology

### Collection of *Norovirus* proteins for vaccine preparation

The three proteins capsid, polyproteins, and small basic proteins were collected from UniProtKB (https://www.uniprot.org) (33). These proteins where found antigenic through VaxiJen server (http://www.ddg-pharmfac.net/vaxijen/VaxiJen/VaxiJen.html) and they were considered for vaccine designing (34-36).

### Prediction of the (CTL) epitope

NetCTL 1.2 is an online server (http://www.cbs.dtu.dk/services/NetCTL/) in which CTL epitopes were predicted against these proteins (37). Prediction of these CTL epitopes is based upon three essential parameters, including peptide attached to MHC-proteasomal C-terminal degradation activity, TAP (Transporter Associated with Antigen Processing) delivery accuracy and I. Artificial neural network was used to predict the attachment of peptide to MHC-I and proteasomal C-terminal degradation while TAP deliverance score was predicted by the weight matrix. For CTL epitope identification threshold was set as 0.75.

### Prediction of helper T-cell epitopes

IEDB (Immune Epitope Database) is an online server (http://tools.iedb.org/mhcii/), which is employed for prediction of Helper T cells lymphocytes of 15-mer length for five human alleles (HLA-DRB1*01:01, HLA-DRB1*01:02, HLA-DRB1*01:03, HLA-DRB1*01:04, HLA-DRB1*01:05) (38-40). This server speculates epitopes based on receptor affinity, which is figured out from IC50 value (binding score) given to each epitope. Epitopes with higher binding affinity mostly have IC50 score <50 nM. The IC50 value determines epitopes affinity, value such is <500 nM represent moderate affinity while <5000 nM instantly show epitopes having low affinity. The affinity of epitopes is inversely related to the value of percentile rank.

### Prediction of B cell lymphocytes

ABCpred is an online server (http://crdd.osdd.net/raghava/abcpred/) employed for the prediction of linear B-cell epitopes. The ABCpred predicts B-cell epitopes (linear) with a precision of 75% (0.49 Sensitivity and 0.75 specificity). Conformational B-cell epitopes prediction is carried out by DiscoTope 2.0 an online server (http://www.cbs.dtu.dk/services/DiscoTope/) (41). The process of computation in this method is based on log-odds ratios (ratio among amino acid composition in epitopes and residues of non-epitopes) and surface accessibility. The sensitivity and specificity are found 0.47 and 0.75, respectively, on the default threshold (3.7).

### Multi-epitopes vaccine sequence construction

High scoring epitopes including other parameters filtered out the final epitopes from CTL and HTL. These selected epitopes were used to construct the final multi-epitopes subunit vaccine. Linker such as AAY and GPGPG interconnect these epitope sand also enhanced effective separation and presentation of epitopes. These linkers have two important roles; first, they assist binding of HLA-II epitopes and also immune processing, and second, they restrain epitopes numbers by specifying cleavage point (42-45). Further, to boost the immune responses an adjuvant (TLR-7 agonist Beta-defensin 3 (Q5U7J2)) was affixed to the vaccine construct N-terminus by EAAAK linker (46).

### Subunit vaccine and interferon

Online server IFNepitope (http://crdd.osdd.net/raghava/ifnepitope/predict.php) assist in designing and predicting the amino acid sequence from proteins, which have the potential to release IFN-gamma from CD4+ T cells. This server helps in the designing of effective and better subunit vaccine by identifying peptide which binds to MHC II and releases IFN-gamma (47)

### Prediction of vaccine allergenicity

For allergenicity prediction of multi-epitope subunit vaccine AlgPred (http://crdd.osdd.net/raghava/algpred/) web tool was used, in which algorithm like (IgEpitope+SVMc+MAST+ARPs BLAST) was applied (48). This hybrid prediction veracity is about 85% at 0.4 thresholds. The server employed six distant routes to act. These approaches abet allergic protein prediction with perfection.

### Vaccine antigenicity prediction

Online server ANTIGENpro (http://scratch.proteomics.ics.uci.edu/) was used for antigenicity prediction (49). Server to predict protein antigenicity used two approaches; primary protein sequence is presented multiple time, and five different machine-learning algorithms were applied to produce the result based on protein microarray data analysis.

### Physiochemical parameters and identification of domain

ProtParam (http://web.expasy.org/protparam/), an online server was used for the prediction of many physiochemical parameters such as theoretical PI, instability index, amino acid composition, in vitro and in vivo half-life, molecular weight, aliphatic index and also the (GRAVY) which is a grand average of hydropathicity (50).

### Prediction of secondary structure

The primary amino-acid sequence was used to predict the secondary structure of Protein by PDBsum. Sequences that shows homology to vaccine protein were selected for structure prediction. Position position-specific iterated (PSI-BLAST) identified homologous residues.

### Prediction of the tertiary structure

For prediction and analysis of protein tertiary structure, an online server Robetta (http://robetta.bakerlab.org) was utilized, which is an automated tool(51). Protein sequences were submitted in FASTA format to predict 3D structure. Models were generated after parsing the structure into respective domains. This model is based either on comparative modelling or de novo structure; for comparative modelling homologs sequence was identified by BLAST, 3D-Jury or FFAS03, and PSI-BLAST, which is then used as a templet. If homologs were not found, de novo structure was generated using Rosetta fragment insertion method.

### Validation of tertiary structure

To validate the predicted model is an essential step, ProSA-web an online server (https://prosa.services.came.sbg.ac.at/prosa.php) was used for validation of tertiary structure. For input structure, overall quality scoring was computed. The predicted protein will have an error when the scoring characteristic was found arbitrary then native proteins. For statistical analysis of non-bonded interaction ERRAT server was used (https://servicesn.mbi.ucla.edu/ERRAT/) (52), and Ramachandran plot analysis was performed on RAMPAGE server (http://mordred.bioc.cam.ac.uk/~rapper/rampage.php) (53). This server utilized PROCHECK principle to validate the structure of protein through Ramachandran plot and separate plots for Proline and Glycine residues.

### Vaccine and TLR-7 Docking

ZDOCK server (http://zdock.umassmed.edu/) was used for the docking of the final vaccine with TLR-7 (54). The ZDOCK webserver produces quick and accurate complexes. Refinement and post-processing of the complexes were subjected to Firedock server(55).

### MD (Molecular dynamic simulation) of Receptor-Vaccine complex

For selected complexes, amber 14 (56) was used to conduct MD simulation. The system was neutralized and solvated with TIP3P water box. Two stages of energy minimization, followed by gentle heating and equilibration, were performed (57). After equilibration, 30ns simulation was conducted. For post-simulation trajectories analysis (RMSD and RMSF), 2.0 ps time scale was used and for trajectory, sampling using CPPTRAJ and PTRAJ(58)implemented in AMBER 14.

### Codon optimization and in silico cloning of vaccine

For effective expression of the vaccine in a host (*E. coli* strain K12) was performed. Reverse translation and codon optimization were performed using JCAT (Java Codon Adaption tool). Norovirus genome expression is distinct from the expression of vector genome, codon optimization ensures maximum expression of vaccine with in the vector. In order to get the desired result, three additional options were selected, such as prokaryote ribosome binding site, restriction enzymes cleavage and rho-independent transcription termination. JCat output includes a codon adaptation index (CAI) and to ensure the high-level protein expression percentage GC content is used (25). In order to clone the desired gene in *E. coli* pET-28a (+) two restriction sites NdeI and XhoI were added to the C and N terminal of the sequence, respectively. Finally, the adapted sequence having the restriction site was inserted to pET-28a (+) to maximize the vaccine expression.

## Acknowledgements

Dong-Qing Wei is supported by the grants from the Key Research Area Grant 2016YFA0501703 of the Ministry of Science and Technology of China, the National Natural Science Foundation of China (Contract no. 61832019, 61503244), the Science and Technology Commission of Shanghai Municipality (Grant: 19430750600), the Natural Science Foundation of Henan Province (162300410060) and Joint Research Funds for Medical and Engineering and Scientific Research at Shanghai Jiao Tong University (YG2017ZD14). The computations were partially performed at the Pengcheng Lab and the Center for High-Performance Computing, Shanghai Jiao Tong University.

